# Development of a Genomic DNA Reference Material Panel for Thalassemia Genetic Testing

**DOI:** 10.1101/676015

**Authors:** Zhen-Zhen Yin, Shou-Fang Qu, Chuan-Feng Huang, Fang Chen, Jian-Biao Li, Shi-Ping Chen, Yu Zheng, Xi Zhang, Xue-Xi Yang, Long-Xu Xie, Ji-Tao Wei, Feng-Xiang Wei, Jian Guo, Jie Huang

## Abstract

Thalassemia is one of the most common autosomal recessive inherited diseases worldwide, and it is also highly prevalent and variable in Southern China. Various types of genetic testing technologies have been developed for diagnosis and screening of thalassemia. Characterized genomic DNA reference materials are necessary for assay development, validation, proficiency testing and quality assurance. However, there is no publicly available reference materials for thalassemia genetic testing as yet. To address the need for these materials, the National Institutes for Food and Drug Control and the China National Gene Bank established 31 new cell lines with 2 wild genotypes and 29 distinct genotypes of thalassemia which account for approximately 90% thalassemia carriers in China. The genomic DNA of 31 cell lines were characterized by four clinical genetic testing laboratories using different genetic testing methods and technology platforms. The genotyping results were concordant among four laboratories. In addition, the results of stability test demonstrated that the genotypes of these DNA samples were not influenced by preanalytical conditions such as long-term exposure to high temperature(37□) environment and repeated freeze-thawing. In conclusion, we developed the first national panel of 31 genomic DNA reference materials which are renewable and publicly available for the quality assurance of various genetic testing methods and will facilitate research and development in thalassemia genetic testing.

## 1 Introduction

As one of the most common monogenic inherited diseases in the world, thalassemia has a high prevalence rate of ranging from 2.5% to 15% in tropical and subtropical areas, including Southern China (1, 2). Thalassemia is caused by genetic defects in human hemoglobin genes, which lead to a decrease or absence of one of the hemoglobin subunits. According to the defective hemoglobin genes, thalassemia is generally classified into two categories: α thalassemia and β thalassemia(3). α thalassemia is frequently caused by deletion, while β thalassemia mainly is caused by single nucleotide variation(SNV) or oligonucleotide insertion/deletion(Indel)(4). Thalassemia is usually inherited in an autosomal recessive pattern and is mainly classified as thalassemia trait, thalassemia intermedia and thalassemia major based on clinical severity. Individuals with thalassemia trait usually have no or mild symptoms of anemia. However, patients with severe α thalassemia (such as hydrops fetalis) often died in utero or shortly after birth, and patients with β thalassemia major require lifelong blood transfusion and special treatment to sustain their life, which brings great burden to their families as well as society(5). Accordingly, screening and accurate diagnosis of thalassemia and prevention of the birth of fetuses with thalassemia major are extremely important. Genetic testing is the gold standard for diagnosis of thalassemia. There are a variety of genetic testing methods for thalassemia diagnosis, such as quantitative polymerase chain reaction(qPCR), PCR-Reverse Dot Blot, PCR-flow through hybridization, target capture sequencing, Gap-PCR, next-generation sequencing(NGS) and Sanger sequencing(6-9). However, most assays for thalassemia diagnosis were laboratory-developed tests and are not approved by the Food and Drug Administration(FDA).

According to the regulatory and accreditation requirements of clinical genetic tesing laboratories, genomic DNA reference materials(RMs) were recommended to use in the assay development, validation and quality control by professional guidelines(10-12). A set of genomic DNA RMs for genetic testing of some disorders and genes, such as Rett Syndrome, Niemann-Pick disease type A, Cystic Fibrosis, familial dysautonomia, HLA loci, BRAC1, BRAC2 and MTHFR, had already been developed(13-17). These well-characterized genomic DNA RMs were usually used to monitor test performance and assure the quality of the genetic testing process in clinical genetic testing laboratories(18-20). Theoretically, genomic DNA RMs should be thoroughly characterized using multiple genetic testing methods and should contain genetic variations common to the corresponding disorder. Therefore, carriers (thalassemia trait, thalassemia intermedia and thalassemia major) derived genomic DNA RMs containing common α- and β-thalassemia variations should be used. It is worth noting that there is no comprehensive characterized genomic DNA RMs panel for thalassemia genetic testing up to now.

For thalassemia, populations in different regions have different spectrums of thalassemia variations (21). There are region-specific types of thalassemia associated variations in China. In order to meet the need for characterized genomic DNA RMs for thalassemia diagnosis in China, the National Institutes for Food and Drug Control(NIFDC) based on the National Key Research and Development Program, in collaboration with China National Gene Bank(CNGB), had characterized the variations of hemoglobin genes in a panel of 31 publicly available genomic DNA samples extracted from cell lines, which were generated by peripheral venous blood samples obtained from normal Chinese individuals and Chinese thalassemia carriers. The publicly available and reproducible characterized genomic DNA RMs help to ensure the accuracy of thalassemia diagnosis and promote research development.

## 2 Materials and Methods

### 2.1 Selection and Collection of Samples

This study was approved by BGI’s institutional review board on bioethics and biosafety and the Ethic Committee of Shenzhen Longgang District Maternity and Child Healthcare Hospital. 31 samples from 29 individuals with α and/or β thalassemia and 2 thalassemia free healthy individuals were selected based on the genotyping information from earlier genetic testing. Written informed consents of all participants were obtained. 5 ml peripheral venous blood was collected in an Heparin lithium vacutainer from all 31 participants.

### 2.2 Cell Line Generation and DNA extraction

Immortalized B lymphoblastoid cell lines were obtained by transformation of B lymphocytes using Epstein-Barr virus as described previously(22, 23). Genomic DNA was extracted from the established immortalized B lymphoblastoid cell lines according to the manufacturer’s instructions of Qiagen DNA extraction kit (Valencia, CA, USA)(24).

### 2.3 Protocol

Four accredited clinical genetic testing laboratories were invited to attend the test for these DNA samples by their current genetic testing methods for thalassemia. The following techniques were used: PCR-flow through hybridization, PCR-Reverse Dot Blot, NGS, and Gap-PCR. A 500ng aliquot of DNA from each cell line was distributed to the clinical genetic testing laboratories for thalassemia genetic testing at the same time. The laboratories were not informed of the expected types of variation in all samples in advance. These samples were genotyped by their standard methods as described blow. Results of these laboratories were sent to the research coordinator (NIDFC), who examined and checked the data for discrepancies. The participating laboratories with discrepancies were required to re-evaluate the relevant samples to determine the cause of the inconsistency without providing the expected genotypes.

#### Laboratory 1(Lab 1)

Testing was performed strictly according to the manufacturer’s protocol of the commercial thalassemia testing kit(Hybribio Limited Corporation, Chaozhou, China)(25). The principle of this kit is based on PCR-flow through hybridization technology which had been approved by the China Food and Drug Administration [REG. NO: CFDA (P) 20153401664](26, 27). 6 types of α thalassemia variations and 16 types of β thalassemia variations were analyzed (Table 1).

**Table 1.** Methods of four laboratories.

| Laboratories | Methods | Detection Range |
| --- | --- | --- |
| Laboratory 1 | PCR-flow through hybridization | $\alpha$: $\alpha^{QS}$, $\alpha^{CS}$, $\alpha^{WS}$, $--^{SEA}$, $-\alpha^{3.7}$, $-\alpha^{4.2}$ |
| | | $\beta$: -29(A>G), -28(A>G), codons 14/15(+G), codon 17(A>T), HbE, codons 27/28(+C), codon 31(-C), codon 43(G>T), codons 41/42(-TCTT), codons 71/72(+A), IVS-I-1 (G>A), IVS-I-1(G>T), IVS-II-654(C>T), IVS-I-5(G>C), Initiation codon(ATG>AGG), 5'UTR; +43 to +40 (-AAAC) |
| Laboratory 2 | PCR-Reverse Dot Blot | $\alpha$: $\alpha^{QS}$, $\alpha^{CS}$, $\alpha^{WS}$, $--^{SEA}$, $-\alpha^{3.7}$, $-\alpha^{4.2}$ |
| | | $\beta$: -28(A>G), -30(T>C), -32(C>A), codons 14/15(+G), codon 17(A>T), HbE, codons 27/28(+C), codon 31(-C), codon 43(G>T), codons 41/42(-TCTT), codons 71/72(+A), IVS-I-1(G>A/T), IVS-I-5(G>C), IVS-II-654(C>T), Initiation codon(ATG>AGG), 5'UTR; +43 to +40(-AAAC) |
| Laboratory 3 | target-NGS | $\alpha$: $\alpha^{QS}$, $\alpha^{CS}$, $\alpha^{WS}$, $--^{SEA}$, $-\alpha^{3.7}$, $-\alpha^{4.2}$, $--^{THAI}$ |
| | | $\beta$: -28(A>G), -29(A>G), -30(T>C), -32(C>A), -90(C>T), -50(G>A), codons 14/15(+G), codon 17(A>T), HbE, codons 27/28(+C), codon 31(-C), codon 43(G>T), Codons 40/41 (+T), codons 41/42(-TCTT), codons 71/72(+A/T), IVS-I-1(G>T), IVS-I-5(G>C), IVS-II-5(G>C), IVS-II-654(C>T), codon 37(G>A), Cap+1(A>C), codon 8(-AA), Initiation codon(ATG>AGG), 5'UTR; +43 to +40(-AAAC), codons 8/9(+AGAA), -10_-7delAACA, HPEH-SEA, $G\gamma(A\gamma\delta\beta)^0$ |
| Laboratory 4 | NGS, Gap-PCR, Sanger sequencing | $\alpha$: non-deletion: all mutations of HBA1 and HBA2<br>deletion: $--^{SEA}$, $-\alpha^{3.7}$, $-\alpha^{4.2}$, $--^{THAI}$, $--^{FIL}$ |
| | | $\beta$: non-deletion: all mutations of HBB<br>deletion: HPEH-SEA, $G\gamma(A\gamma\delta\beta)^0$ |

#### Laboratory 2(Lab 2)

Testing was performed strictly according to the instruction of their own kit for thalassemia (Yaneng Biological technology Co., Ltd., Shenzhen, China). 3 common α thalassemia deletions, 3 non-deletion α thalassemia mutation and 17 β thalassemia variations in Chinese were analyzed using PCR-reverse dot hybridization technique(Table 1).

#### Laboratory 3(Lab 3)

NGS was used for thalassemia genetic testing by this laboratory. Primers were designed according to the reference sequences of the HBA2 and HBB gene (accession numbers NC_000016.9 [222846…223709] and NC_000011.9 [5246696…5248301]) from the NCBI database (http://www.ncbi.nlm.nih.gov/nuccore/NC_000016.9 and http://www.ncbi.nlm.nih.gov/nuccore/NC_000011.9) with the primer premier5. Primers:

SF: GAGGGAAGGAGGGGAGAAGCTGAGT SR: AGCCCACGTTGTGTTCATGGC

3F: CCCCACATCCCCTCACCTACATTC 3R: GCATCCTCAAAGCACTCTAGGGTCC

4F: TGCTTTTGTGAGTGCTGTGTTGACC 4R: GAAGTAGCTCCGACCAGCTTAGCAAT

αF: CCCCACATCCCCTCACCTACATTC αR: CGGGCAGGAGGAACGGCTAC

TF: GACCATTCCTCAGCGTGGGTG TR: CAAGTGGGCTGAGCCCTTGAG

BF: CAGAAGAGCCAAGGACAGGTACGGCT

BR: AAGGGCCTAGCTTGGACTCAGAATAATCC

HF: CCAGCCTCATGGTAGCAGAATC HR: TGGTATCTGCAGCAGTTGCC

GF: TCAACAATTATCAACATTACAC GR: GGCATATATTGGCTCAGTCA

The PCR products were used to construct the library by the Ion Xpress™ Plus Fragment Library Kit (Life Technologies, Carlsbad, USA). Each library was diluted to 200 pM according to its quantified concentration as determined on the Qubit 3.0 fluorometer. Thirty-two libraries of 100 pM were mixed together and emulsion PCR amplified on the Ion PI™ Hi-Q™ ISPs using the Ion OneTouch™ 2 Instrument (Life Technologies, Carlsbad, USA) according to the manufacturer’s instructions. The template-positive ISPs were enriched and loaded onto one Ion PI chip and sequenced on the Ion Torrent Proton Machine (Life Technologies, Carlsbad, USA) which is a semiconductor sequencing platform. This method can detect 3 common α thalassemia gene mutations, 4 α thalassemia deletions, 27 β thalassemia mutations and 2 rare β thalassemia deletions in Chinese (Table 1).

#### Laboratory 4(Lab 4)

Combined NGS and Gap-PCR was used for thalassemia genotyping by Lab 4. Point mutations were detected by NGS. 5 α thalassemia deletions and 2 rare β-golbin gene deletions were analyzed by Gap-PCR(Table 1). The detailed protocol of characterizing thalassemia associated variations was described previously(28, 29).

### 2.4 Stability test

Each of the thirty-one DNA samples from cell lines were aliquoted into four portions. Two portions were incubated at 37□ for one month and another two portions were subjected to 4 repeated freeze-thawing cycles every five days until one month. Afterwards, the hemoglobin genotypes of these samples were reanalyzed. Point mutations were analyzed by NGS and validated by Sanger sequencing, while copy number variations were detected by Gap-PCR.

## 3 Results

### 3.1 Establishment of cell lines

Two samples with wild genotype and twenty-nine samples containing 25 different variations with 29 distinct genotypes were selected to establish cell lines for the first national genomic DNA RMs panel for thalassemia genetic testing(Table 2). The 18 most common hemoglobin genotypes are as follows: --^SEA^/αα, -α^4.2^/αα, -α^3.7^/αα, α^WS^α/αα, α^CS^α/αα and α^QS^α/αα, Codons 41/42 (-TTCT)/β^N^, Codon 17 (A>T)/β^N^, −28 (A>G)/β^N^, Hb E/β^N^, −50(G>A)/β^N^, Initiation codon (ATG>AGG)/β^N^, Codons 71/72 (+A)/β^N^, IVS-I-1(G>T)/β^N^, Codon 43(G>T)/β^N^, Codons 27/28(+C)/β^N^, −29 (A>G)/β^N^, Codons 14/15(+G)/β^N^. Six rare hemoglobin genotypes were also included: −90 (C>T)/β^N^, 5’UTR; +43 to +40 (-AAAC)/β^N^, Codon 37(G>A)/β^N^, SEA-HPFH/β^N^, Gγ(Aγδβ)^0^/β^N^ and --^THAI^/αα. In addition, five composite genotypes, -α^3.7^/--^SEA^, Codons 41/42 (-TTCT)/Gγ(Aγδβ)^0^, IVS-II-654 (C>T)/β^N^+--^SEA^/αα, IVS-II-654 (C>T)/IVS-II-654 (C>T)+--^SEA^/αα, and -α^4.2^/-α^4.2^+IVS-II-654/β^N^ were also included. Thirty-one immortalized B lymphoblastoid cell lines were all successfully generated.

**Table 2.** The hemoglobin genotypes of 31 samples and the genetic testing results of four laboratories.

| Samples | Genotypes | Lab 1 | Lab 2 | Lab 3 | Lab 4 |
| --- | --- | --- | --- | --- | --- |
| 1 | Wild type | √ | √ | √ | √ |
| 2 | Hb E/$\beta^N$ | √ | √ | √ | √ |
| 3 | -28 (A>G)/$\beta^N$ | √ | √ | √ | √ |
| 4 | IVS-II-654 (C>T)/$\beta^N$ | √ | √ | √ | √ |
|  | --<sup>SEA</sup>/αα |  |  |  |  |
| 5 | --<sup>SEA</sup>/αα | √ | √ | √ | √ |
| 6 | -α<sup>3.7</sup>/--<sup>SEA</sup> | √ | √ | √ | √ |
| 7 | -α<sup>3.7</sup>/αα | √ | √ | √ | √ |
| 8 | IVS-I-1 (G>T)/β<sup>N</sup> | √ | √ | √ | √ |
| 9 | Wild type | √ | √ | √ | √ |
| 10 | Codons 71/72 (+A)/β<sup>N</sup> | √ | √ | √ | √ |
| 11 | Codon 17 (A>T)/β<sup>N</sup> | √ | √ | √ | √ |
| 12 | α<sup>QS</sup>α/αα | √ | √ | √ | √ |
| 13 | 5'UTR; +43 to +40<br>(-AAAC)/β<sup>N</sup> | √ | √ | √ | √ |
| 14 | Codon 43 (G>T)/β<sup>N</sup> | √ | √ | √ | √ |
| 15 | Initiation codon<br>(ATG>AGG)/β<sup>N</sup> | √ | √ | √ | √ |
| 16 | Codon 37(G>A)/β<sup>N</sup> | Out of<br>detection<br>range | Out of<br>detection<br>range | √ | √ |
| 17 | Codons 41/42<br>(-TTCT)/β<sup>N</sup> | √ | √ | √ | √ |
| 18 | -α<sup>4.2</sup>/αα | √ | √ | √ | √ |
| 19 | Codons 27/28 (+C)/β<sup>N</sup> | √ | √ | √ | √ |
| 20 | α<sup>CS</sup>α/αα | √ | √ | √ | √ |
| 21 | α<sup>WS</sup>α/αα | √ | √ | √ | √ |
| 22 | -50 G>A/β<sup>N</sup> | Out of<br>detection<br>range | Out of<br>detection<br>range | √ | √ |
| 23 | -29 (A>G)/β<sup>N</sup> | √ | √ | √ | √ |
| 24 | Gγ(Aγδβ)<sup>0</sup>/β<sup>N</sup> | Out of<br>detection<br>range | Out of<br>detection<br>range | √ | √ |
| 25 | Codons 41/42<br>(-TTCT)/Gγ(Aγδβ)<sup>0</sup> | Out of<br>detection<br>range | Out of<br>detection<br>range | √ | √ |
| 26 | Codons 14/15 (+G)/β<sup>N</sup> | √ | √ | √ | √ |
| 27 | -90 (C>T)/β<sup>N</sup> | Out of<br>detection<br>range | Out of<br>detection<br>range | √ | √ |
| 28 | -α<sup>4.2</sup>/-α<sup>4.2</sup><br>IVS-II-654/β<sup>N</sup> | √ | √ | √ | √ |
| 29 | IVS-II-654<br>(C>T)/IVS-II-654 (C>T)<br>--<sup>SEA</sup>/αα | √ | √ | √ | √ |
| 30 | SEA-HPFH/β<sup>N</sup> | Out of | Out of | √ | √ |
|  |  | detection<br>range | detection<br>range |  |  |
| 31 | --<sup>THAI</sup>/$\alpha\alpha$ | Out of<br>detection<br>range | Out of<br>detection<br>range | √ | √ |

√ indicates expected mutation is detected by the method; × indicates the expected mutation is not detected by method.

### 3.2 Characterization of genomic DNA samples

DNA Samples from the 31 cell lines were prepared for genetic testing in four laboratories. And the hemoglobin genotypes of these DNA samples were characterized by four genetic testing methods used by the laboratories. The detection ranges of these methods are summarized in Table 1. The results of two laboratories (Lab 3 and Lab 4) showed 100% concordance among all 31 samples. The hemoglobin genotypes were totally consistent with expectations. The other two laboratories (Lab 1 and Lab 2) also got the expected genotypes of 24/31 samples, while the genotypes of the remaining 7/31 samples (No.16, 22, 24, 25, 27, 30, 31) were not identified correctly because their genotypes are out of the detection range. In general, the expected hemoglobin genotypes of all DNA samples were confirmed by four laboratories and assay platforms. Therefore, all 31 genomic DNA samples were characterized as candidate thalassemia genomic DNA RMs.

### 3.3 Stability test of 31 DNA samples

In the stability test, we investigated the influence of long-term exposure to high temperature(37□) and repeated freeze-thawing. Two portions of candidate 31 genomic DNA RMs were incubated at 37□ for one month and were subjected to 4 repeated freeze-thawing cycles every five days until one month, respectively. The reanalyzed results of hemoglobin genotypes of these treated DNA samples were fully consistent with expected genotypes(data not shown), which demonstrated an excellent stability of the candidate 31 genomic DNA RMs.

## 4 Disscusion

This study describes the establishment, characterization and stability test of candidate 31 genomic DNA RMs for thalassemia genetic testing, which could provide an inexhaustible genetic resources for hemoglobin genotyping. Two samples with wild genotypes and twenty-nine samples containing 25 different variations with 29 distinct genotypes were selected. Six common variations and one rare Thailand deletion of α thalassemia, which account for 99.61% of α thalassemia carriers, were included(30, 31). The Thailand deletion has an allele frequency of about 0.05% in mainland China and was previously reported in Taiwan aboriginals and Southeast Asian(32, 33). 13 common and 5 rare β thalassemia variations in China were also included. In addition, four invaluable samples with composite hemoglobin genotypes were obtained from four thalassemia major patients. The variations and genotypes of hemoglobin genes of 29 thalassemia-positive samples represent approximately 90% thalassemia carriers in China. Furthermore, another two thalassemia free samples as negative control were also included. The genotyping results of all 31 samples were concordant among four laboratories and technology platforms and showed an outstanding stability among different preanalytical conditions.

Thalassemia is a popular monogenic disease in tropical and subtropical regions. There is a high prevalence rate of thalassemia in Southern China. Prenatal diagnosis is the only effective way to prevent the birth of fetuses with severe thalassemia. Genetic testing is critical for prenatal diagnosis or preimplantation genetic diagnosis for at risk couples with thalassemia. With the rapid progress of genetic testing technologies in recent years, a number of genetic testing techniques had been used for thalassemia genotyping, only a few of which had been approved by FDA. Well characterized genomic DNA RMs has been recognized as a vital role in the molecular genetic testing for disease control and prevention(34, 35). Genomic DNA RMs were usually used to help the design of assay, and the establishment of performance characteristics such as precision, accuracy, analytical sensitivity and specificity.

However, none commercial genomic DNA RMs for thalassemia genetic testing was available as yet. In the absence of publicly available genomic DNA RMs for thalassemia, the majority of thalassemia genetic testing laboratories had to use residual available DNA from patients specimens for quality control. However, residual specimens are generally nonrenewable and limited in supply, and they may not be completely characterized and have poor stability. For example, it is possible that the residual specimen was contaminated or degraded so that the actual hemoglobin genotype is different from the genotype provided. Furthermore, as the need of quality assessment of thalassemia assays, as many variations as possible in the populations should be included in the genomic DNA RMs. However, residual specimens with various kinds of variations, which could represent the spectrum of thalassemia variations that occur in thalassemia carriers and patients, are difficult to obtained.

The genomic DNA RMs panel for thalassemia in this study contained a varity of common and rare variations that represents the vast majority of thalassemia carriers in China. The characterized genomic DNA samples in this study could help to solve these problems and provide a publicly available and limitless source for clinical thalassemia genetic testing. The establishment of this relatively comprehensive genomic DNA reference material panel is of great significance for assay development, validation, quality control, proficiency testing and other research applications in thalassemia. These genomic DNA RMs are publicly available from the NIFDC(http://www.nifdc.org.cn/bzwz/CL0481/). More information about the genomic DNA RMs and cell lines are available at the NIFDC website.

